# Realfreq: Real-time base modification analysis for nanopore sequencing

**DOI:** 10.1101/2025.01.23.634192

**Authors:** Suneth Samarasinghe, Ira Deveson, Hasindu Gamaarachchi

**Affiliations:** School of Computer Science and Engineering, UNSW Sydney, 2052, NSW, Australia; Genomics and Inherited Disease Program, Garvan Institute of Medical Research, 2010, NSW, Australia; Centre for Population Genomics, Garvan Institute of Medical Research and Murdoch Children’s Research Institute, 2010, NSW, Australia; St Vincent’s Clinical School, UNSW Sydney, 2052, NSW, Australia

**Keywords:** Real-time, Base modifications, Methylation, Nanopore

## Abstract

**Motivation:** Nanopore sequencers allow sequencing data to be accessed in real-time. This allows live analysis to be performed, while the sequencing is running, reducing the turnaround time of the results. We introduce *realfreq*, a framework for obtaining real-time base modification frequencies while a nanopore sequencer is in operation.

**Results:** *Realfreq* calculates and allows access to the real-time base modification frequency results while the sequencer is running. We demonstrate that the data analysis rate with *realfreq* on a laptop computer can keep up with the output data rate of a nanopore MinION sequencer, while a desktop computer can keep up with a single PromethION 2 solo flowcell.

**Availability and Implementation:** Realfreq is a free and open-source application implemented in C programming language and shell scripts. The source code and the documentation for *realfreq* can be found at https://github.com/imsuneth/realfreq.

## Introduction

Nanopore sequencing has become increasingly popular with its ability to sequence long reads of both DNA and direct-RNA strands while providing real-time modification data alongside basecalled data (Marx [2023], Wang et al. [2021], Jain et al. [2022], Xu and Seki [2020]). Nanopore sequencers measure ionic currents when DNA or RNA strands pass through nanopores, and this current signal data can be accessed in real-time during the sequencing process. Such real-time access enables adaptive sampling, that can reject unwanted molecules during sequencing, and supports live data analysis (e.g., live basecalling, live modification calling), reducing the turnaround time of results (Kovaka et al. [2021], Payne et al. [2021], Sneddon et al. [2024]). These adaptive sampling and live analysis methods now even come built into the MinKNOW sequencing acquisition software from Oxford Nanopore Technologies which manufactures nanopore sequencers (Wang et al. [2024]).

However, the live analysis of nanopore data is currently limited to basecalling and modification calling followed by sequence alignment, operations that can be independently performed on individual reads. Further downstream analysis, such as variant calling and calculating modification frequencies, are still performed post-hoc, once the sequencing run is complete and all the sequenced reads are available. Methods to perform the complete analysis workflow in real-time would be beneficial for fully unlocking the potential of nanopore sequencing, further reducing the time to results. The urgency in results applies to time-critical contexts such as clinical diagnostics, where rapid identification of genetic and epigenetic markers can guide immediate treatment decisions (Hench et al. [2024]). Another application would be in forensics, where rapid identification of methylation patterns can offer valuable clues in criminal investigations and reduce the time required to solve cases (Yuen et al. [2024]).

We present *realfreq*, a framework that enables the real-time computation of base modification frequencies while the nanopore sequencer is in operation. *Realfreq* watches the raw signal files (e.g., POD5 files) written by the nanopore sequencer onto the host computer’s disk, processes them, and provides base modification frequencies in real-time. *Realfreq* periodically writes the modification frequencies to the disk and also at the end of the sequencing run, making the results available during sequencing and as soon after completion. *Realfreq* can recover and resume operation in the event of a host system crash. *Realfreq* also offer methods for third-party applications to utilise these real-time modification frequencies using a server-client configuration.

### Real-freq Framework

*Realfreq* framework consists of three main components: the *realfreq-monitor* that watches the sequencing experiment directory, *realfreq-pipeline* that is a series of tools performing up to the alignment step, and the *realfreq-program* that performs the real-time modification frequency computation (Fig. 1a). This modular design in *realfreq* makes it possible to support a wide range of analysis workflows. The modularity also offers users the flexibility to either utilise the entire *realfreq* framework or selectively use its individual components, depending on their needs.

**Fig. 1:**
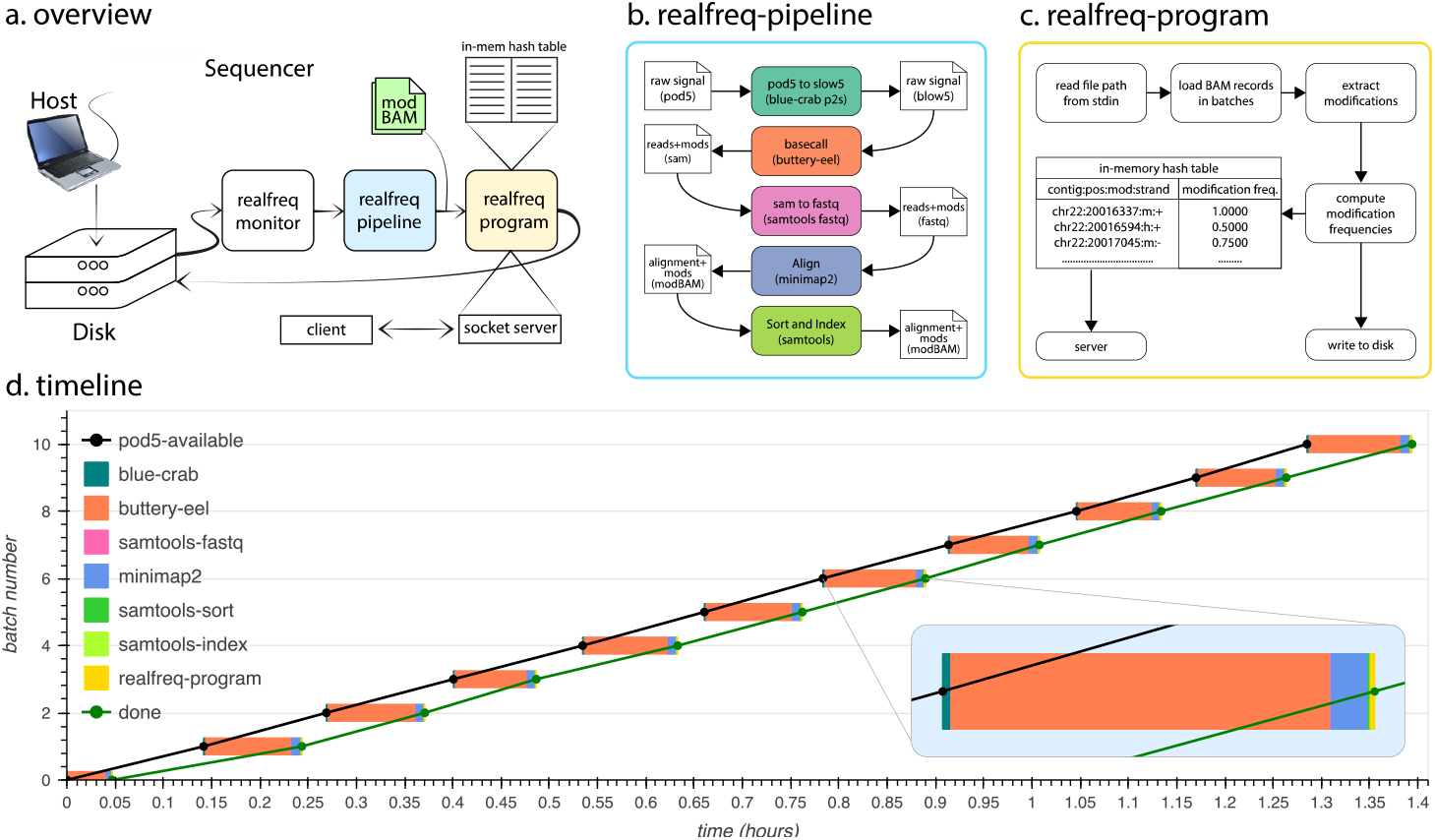
**a)** Overview of *realfreq*. **b)** Data flow and tools used in the *realfreq-pipeline* for processing raw signal files into modBAM. **c)** Structure of *realfreq-program* that reads modBAMs, computes and updates modification frequencies in the in-memory hash table. **d)** A timeline diagram when executing *realfreq* on a laptop for a MinION sequencing run. x-axis (up to 1.5 hours of the sequencing run) is the time and y-axis is the batch number (11 raw signal batches, so 0 to 10). Black and green dots represent the availability of POD5 and completion of realfreq modification frequency computation for each batch, respectively. Black and green lines drawn connecting black and green dots are parallel to each other throughout the sequencing run, demonstrating that realfreq can keep up processing with the output rate of sequencing. The breakdown of time taken for different processing steps in the pipeline (steps in b) for each data batch is shown using coloured boxes, with one batch zoomed in.

#### Realfreq-monitor

The *realfreq-monitor* watches the sequencing experiment directory on the host computer where MinKNOW writes files containing batches of reads. This experiment directory (e.g. */data/exp_id* or */var/lib/minknow/data/exp_id*) can be specified by the user. By default, *realfreq-monitor* watches for newly created POD5 files containing raw signal data. As soon as a new POD5 file is found, the *realfreq-monitor* triggers the execution of the *realfreq-pipeline* (see below) on that POD5 file. If MinKNOW has been configured to perform live modification calling and alignment, the user could alternatively specify *realfreq-monitor* to watch for newly created aligned BAM files containing modification tags (referred to as modBAM).

#### Realfreq-pipeline

*Realfreq-pipeline* is a shell script that invokes a series of tools to generate alignment files (BAM) containing base modification data (Fig. 1b), starting from POD5 files. By default, the *realfreq-Pipeline* converts POD5 to BLOW5 using *blue-crab* (Gamaarachchi et al. [2022]); performs base and modification calling using *buttery-eel* (Samarakoon et al. [2023]) through ONT Dorado server; converts SAM to FASTQ using samtools (Danecek et al. [2021]); and, aligns reads using *minimap2* (Li [2018]). Since this pipeline is implemented as a shell script, users can easily modify specific commands/parameters or fully modify the workflow to incorporate the tools of their choice. The user can also choose to completely bypass the *realfreq-pipeline* and simply use the BAM files created by the MinKNOW’s inbuilt live modification calling feature, if they wish.

#### Realfreq-program

*Realfreq-program* forms the core of *realfreq*. It is an efficient modification frequency computation tool developed to support a continuous input data stream from a nanopore sequencer. *Realfreq-program* performs modification calling based on base-modification tags embedded in the modBAM file. *Realfreq-program* opens and reads a modBAM file path specified from the *stdin* and updates an in-memory hash table of modification frequencies (Fig. 1c). This process is repeated for the next modBAM file path on the *stdin*.

*Realfreq-program* periodically writes the modifications frequency table to disk in bedmethyl or TSV format. *Realfreq-program* also has the *realfreq-server* built into it providing an interface for third-party applications to retrieve base modification information in real-time.

### Usage

*Realfreq* can be launched on the host computer that runs MinKNOW by calling the parent script *realfreq*.*sh*. The user can point the script with the sequencing experiment directory and provide the command line arguments as follows:

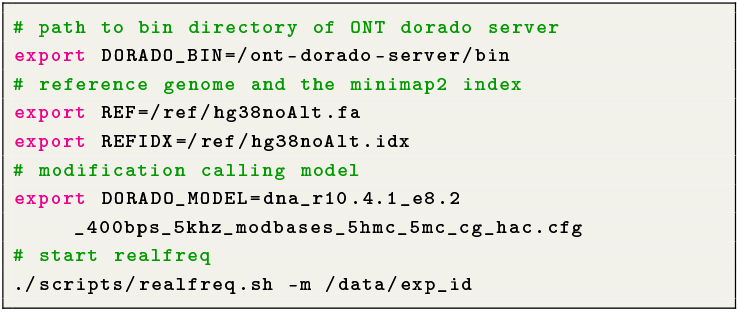

The above example uses the default *realfreq-pipeline* that doesn’t rely on MinKNOW live analysis. If live modification calling and alignment are enabled in MinKNOW, the user can specify the *-a bam* option to monitor the modBAM files directly written to the experiment directory (Supplementary Note 1). *Realfreq* also allows the user to recover and resume analysis in case of a host system crash, which can be invoked with the *–r* flag (Supplementary Note 1).

*Realfreq* periodically updates the modification frequency output (TSV by default, BED if-b is specified) on the disk in the experiment directory. By default, this is per each input file. A time interval can be given via-w argument (Supplementary Note 1). The periodic updates to the modification frequency output during the sequencing allow the user or third-party programs to probe the file for accessing the data periodically, for instance for those who want to access every hour or so.

A third-party application that would require the most up-to-date frequency results and would want to access them very frequently can use the server-client method. In the server-client method (activated via option *–c PORT*, Supplementary Note 2), *realfreq* acts as the server and the user application as the client. It provides an interface to query modification frequencies using simple socket connections. Supported query formats and querying methods can be found in Supplementary Note 2. This interface could be used by dynamic adaptive sampling methods (Weilguny et al. [2023]) to perform modification frequency-aware selective sequencing. For instance, an application may check if the coverage of reads with high-confident modification calls within a genomic region or site of interest has reached an adequate threshold and then reject reads for such regions.

*Realfreq-program* can be alternatively used as a standalone program and thus can be integrated into an existing workflow to compute live modification frequency. The file path input can be either piped or given as a list through standard input as described in Supplementary Note 3.

## Evaluation

*Realfreq* was first evaluated on a laptop computer connected to a MinION (Methods and Implementation). *Realfreq* could keep up the processing pace with the data generation, that is, a batch of data (POD5 file with 4000 reads) could be processed before the next batch of data was available (Fig. 1d). Processing took ∼75% of the time between two data batches. The *realfreq-program* that computes the modification frequency only took ∼1.1% out of the total time for processing a batch. The majority of the processing time was spent on base+modification calling (∼87.3%), followed by ∼8.7% for the alignment and ∼2.4% for data conversion. The peak RAM usage of *realfreq-program* was recorded at 5.8GB throughout the test. Base+modification calling was performed using the high accuracy model. We also tested *realfreq* an alternate pipeline where *f5c* (Gamaarachchi et al. [2020]) is used for methylation calling. The processing could still keep up with the data generation (Supplementary Fig. S1).

Furthermore, we tested on a small desktop computer connected to a Promethion 2 solo for a sequencing run of 48 hours (Methods and Implementation). Real-freq could keep up the processing with the data rate of the sequencer (Supplementary Fig S2), with processing (from base+modification calling to *realfreq-program*) taking ∼81% of the time between two data batches. The peak RAM usage of *realfreq-program* was recorded at 9.4 GB throughout the test. Base+modification calling was performed using the high accuracy model.

While the above experiments were on DNA for 5mC methylation (R10.4.1 chemistry), realfreq was also evaluated on RNA for modifications (m6A calling using super accuracy model, RNA004 chemistry), which kept up the processing pace Supplementary Fig. S3).

## Methods and Implementation

### Realfreq-monitor and realfreq-pipeline scripts

*Realfreq-monitor* uses the inotify API which is a Linux kernel subsystem built on top of the Virtual File System (VFS) layer to monitor file system events, hence avoiding overheads of polling methods to track newly created files. Once a new raw signal file is created, the *realfreq-monitor* outputs its absolute path to standard output. The *realfreq-monitor* is adapted from real-time FAST5 to BLOW5 conversion scripts (Gamaarachchi et al. [2022]).

When inotify detects a new file, *realfreq-monitor* ensures that the file type is correct and it has not already been processed before writing the file path to standard output that triggers *realfreq-pipeline*.

### Realfreq-program

*Realfreq-program* is implemented using C programming language. It takes the reference genome as an argument and keeps scanning the standard input. When a modBAM file path is found, it loads the modBAM, extracts the modifications, computes modification frequencies, and updates the results in a hash table. *Realfreq-program* terminates when the standard input reaches the end of file (eof).

We use [htslib](https://github.com/samtools/htslib) to load modBAM files and to retrieve MM (modification information) and ML (modification probabilities) base modification tags that are described in [SAM tags specification](https://samtools.github.io/hts-specs/SAMtags.pdf). Then, we decode MM and ML tags, extract modifications of interest, and update modification frequencies on an in-memory hash table implemented using *khash* from [klib](https://github.com/attractivechaos/klib), a lightweight hash table with a small memory footprint and overhead. Decoding tags and extracting modification are performed using multiple threads implented via POSIX threads. Updating of the hash table is performed using a single thread to avoid multiple threads from being locked.

*Realfreq-program* periodically writes the hash table onto the disk as a binary file and a list of paths of processed files as a log file. The binary file and the processed file list is used to resume in case of failure. When resuming, *realfreq-program* loads the hash table (binary file), while *realfreq-monitor* uses the list of processed file to avoid any reprocessing.

The server-client interface in *realfreq-program* is a TCP socket server that can handle multiple client connections.

### Experiment setup

The laptop computer (used for the evaluation of the DNA methylation on MinION) was equipped with an Intel i7-1370P CPU with 14 cores/28 threads, 32 GB of RAM, 1TB NVMe SSD storage and an NVIDIA RTX A500 GPU. The laptop was running Ubuntu 22.04 as the O/S and MinKnow version 24.06.14.

The desktop computer (used for DNA methylation on Promethion 2 solo and RNA m6A on MinION) was equipped with an Intel i7-11700F CPU with 8 cores/16 threads, 64 GB of RAM, 4TB of SATA SSD storage, NVIDIA GeForce 3090 GPU and was running PopOS 20.04 as the O/S and Minknow version 23.11.4.

A simulated playback was performed using bulkfast5 files (Data and code availability). See Supplementary Note 4 for more details including commands used and their versions.

## Data and code availability

The source code for realfreq is open-source at https://github.com/imsuneth/realfreq under the MIT license. Bulk-fast5 file for MinION DNA simulation was downloaded from the [publicly available link](https://s3.amazonaws.com/nanopore-human-wgs/bulkfile/GXB02001_20230509_1250_FAW79338_X3_sequencing_run_NA12878_B1_19382aa5_ef4362cd.fast5) provided under the readfish GitHub repository. The bulk-fast5 file for the MinION RNA simulation is available from ENA accession ERR13928856. The bulk-fast5 file for the PromethION DNA simulation is too large (*>*3TB) to be uploaded currently to a public respository.

## Acknowledgement

We thank Igor Stevanovski and Jillian Hammond for generating bulk-fast5 files for the RNA MinION and DNA PromethION runs.

---

**Supplementary Fig. S 1:**
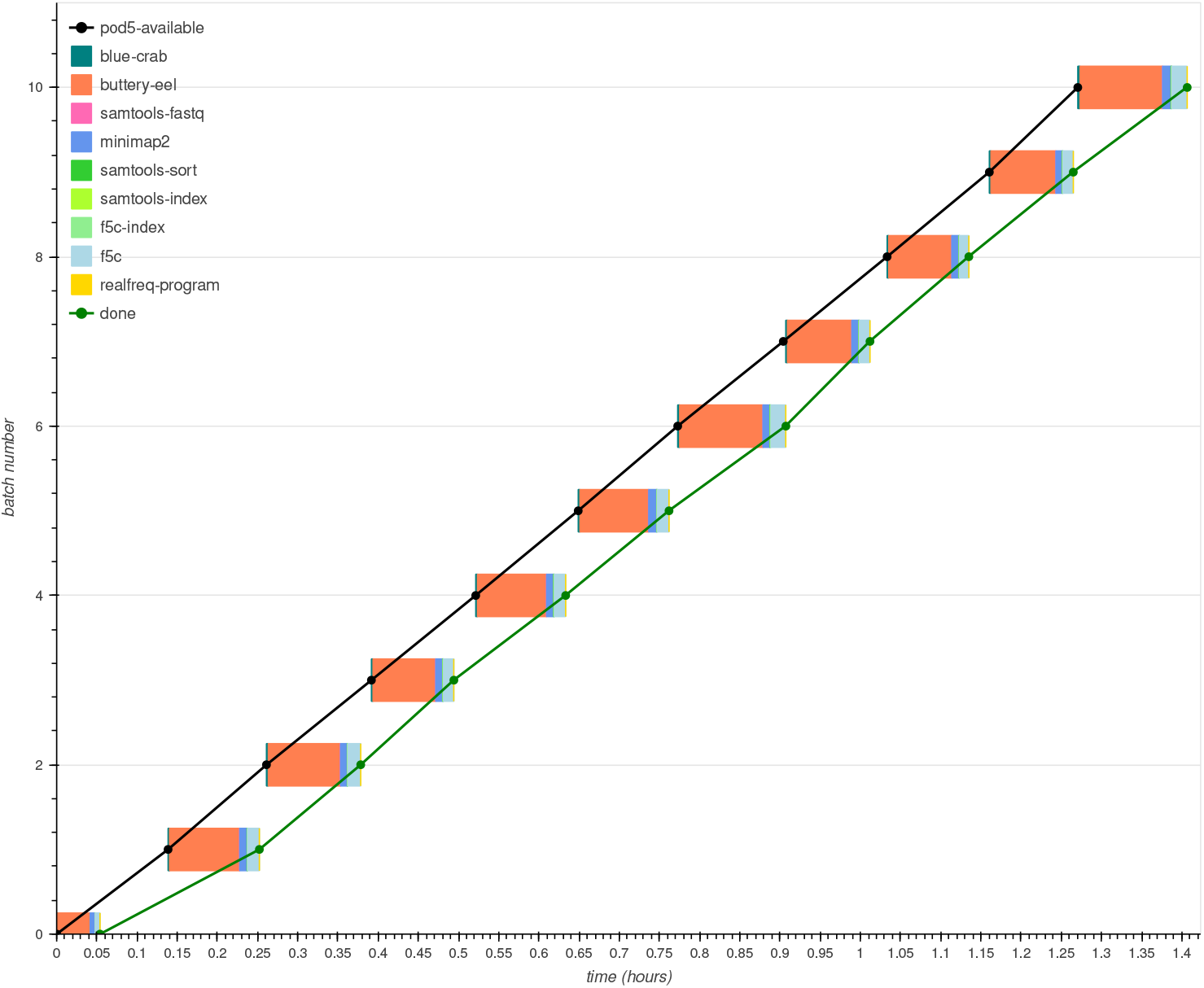
A timeline when executing *realfreq* with the f5c pipeline. This experiment is very similar to Fig. 1d, except that f5c is now used for methylation calling instead of methylation calling within buttery-eel. Black and green dots represent the availability of POD5 and completion of modification frequency computation respectively. Black and green lines connecting black and green dot markers stay parallel throughout the sequencing run, demonstrating that realfreq with f5c also can keep up with the output rate of sequencing.

**Supplementary Fig. S 2:**
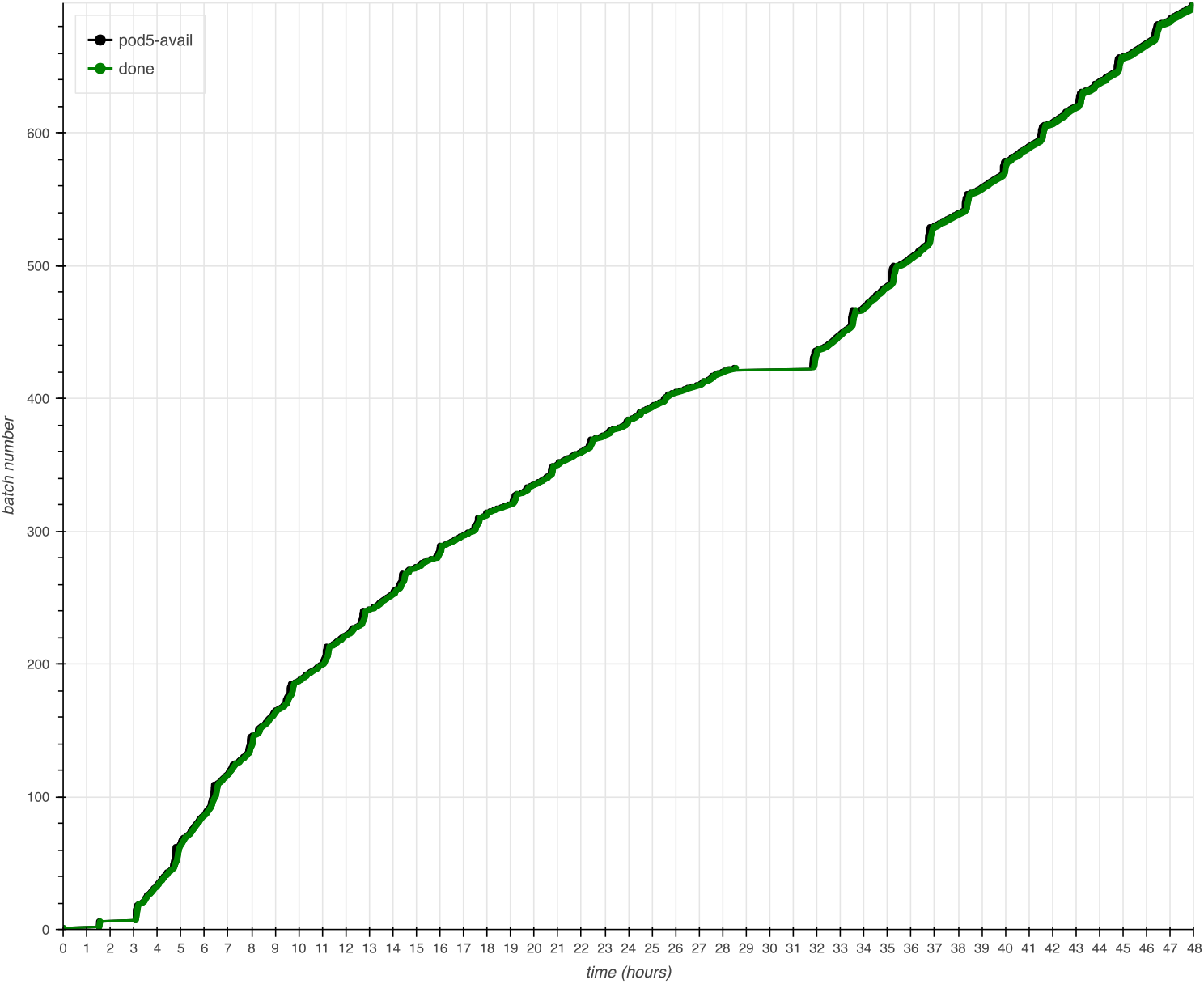
Timeline when executing *realfreq* (default pipeline) on a desktop computer with a PromethION 2 solo single flowcell for a sequencing run of 48 hours. During the sequencing run, ∼700 raw signal files were generated, each containing 4000. Black and green cross dots represent the availability of POD5 and completion of modification frequency computation, respectively. Black and green lines connecting black and green dots are parallel to each other throughout the sequencing run. That demonstrates that *realfreq* can keep up with the output rate of sequencing. Base+modification calling was performed using the high accuracy model.

**Supplementary Fig. S 3:**
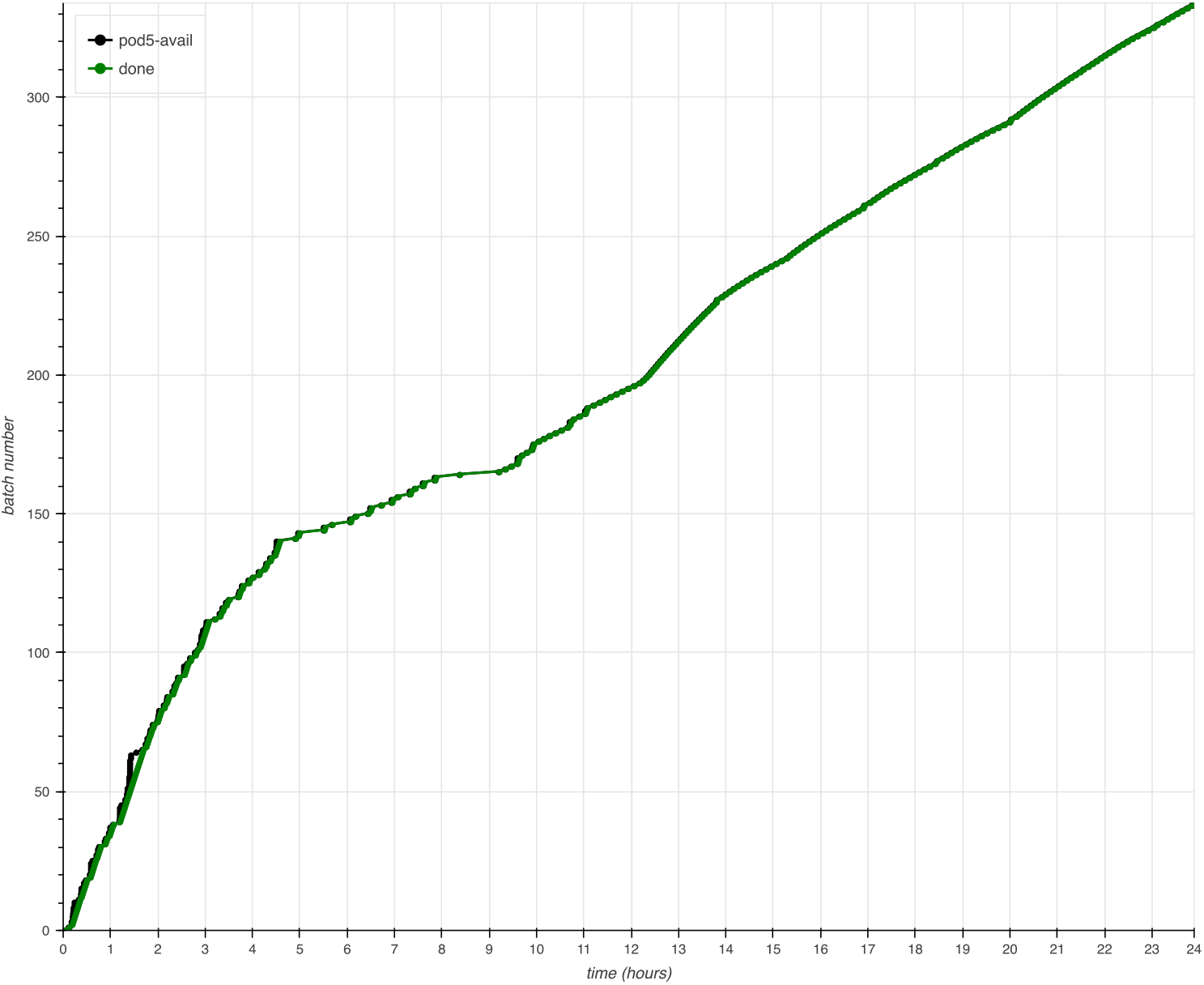
The timeline when executing *realfreq* on a desktop for computing RNA m6A of MinION sequencing run of 24 hours. Around ∼325 raw signal files were generated each containing 4000 reads. Black and green dots represent the availability of POD5 and completion of modification frequency computation respectively. Black and green lines connecting black and green dot markers stay parallel throughout the sequencing run. That indicates realfreq can keep up with the output rate of sequencing. Base+modification calling was performed using the super accuracy model.

