## Supplementary Notes for "Realfreq: Real-time base modification analysis for nanopore sequencing"

### Supplementary Note 1 - Detailed usage of *realfreq*

#### Installation

1. Build
```
sudo apt-get install zlib1g-dev # zlib development files
git clone https://github.com/imsuneth/realfreq
cd realfreq
./scripts/install-hts.sh # htslib
make
```

2. Add to PATH by appending the following line to the ~/.bashrc file.
```
export PATH=$PATH:/parent/dir/of/realfreq
```

3. The following tools should be installed and available to use the default *realfreq-pipeline*.
    - Blue-crab: https://github.com/Psy-Fer/blue-crab
    - Buttery-eel: https://github.com/Psy-Fer/buttery-eel
    - samtools: https://www.htslib.org/download
    - Minimap2: https://github.com/lh3/minimap2

#### Running the default *realfreq-pipeline*

As shown in the figure below, *realfreq-monitor* monitors POD5 files generated by the sequencer. The default *realfreq-pipeline* (pink block in the figure) processes those POD5 files to generate modBAM files. *Realfreq-program* calculates the modification frequencies.

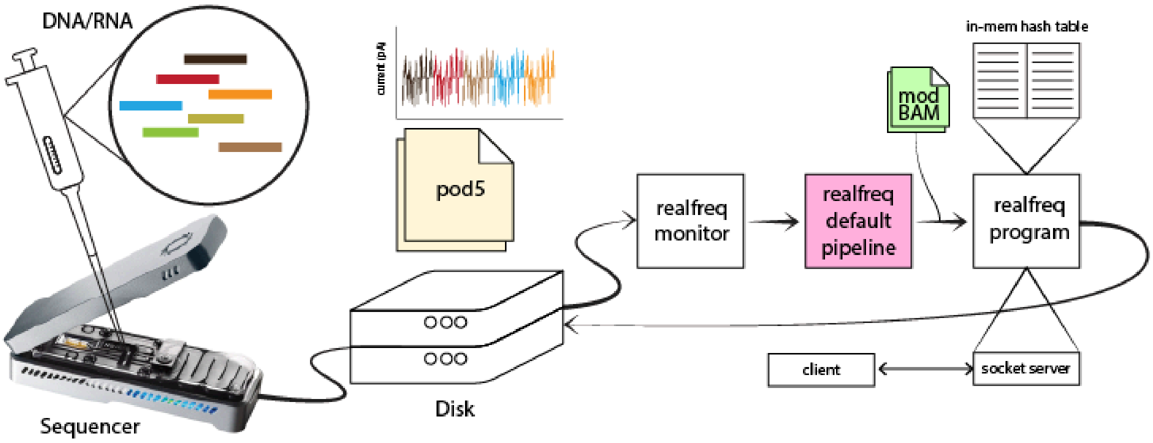


```
export DORADO_BIN=/ont-dorado-server/bin
export REF=/ref/hg38noAlt.fa
export REFIDX=/ref/hg38noAlt.idx
export DORADO_MODEL="dna_r10.4.1_e8.2_400bps_5khz_modbases_5hmc_5mc_cg_hac.cfg"
./scripts/realfreq.sh -m /data/exp_id
```

#### Resuming

*Realfreq-program* keeps a log of processed file paths and periodically writes the current hash table in memory to the disk to be used in case of a failure. When *realfreq.sh* is executed with *-r* option, *realfreq-program* loads the last written hash table from disk to memory and continues processing new files. The missed files are processed at the end of the sequencing run.

```
export DORADO_BIN=/ont-dorado-server/bin
export REF=/ref/hg38noAlt.fa
export REFIDX=/ref/hg38noAlt.idx
export DORADO_MODEL="dna_r10.4.1_e8.2_400bps_5khz_modbases_5hmc_5mc_cg_hac.cfg"
./scripts/realfreq.sh -r -m /data/exp_id
```

#### Running a custom modification calling pipeline

As shown in the figure below, *realfreq-monitor* monitors POD5 files generated by the sequencer. The *realfreq-f5c-pipeline* (light red block in the figure) processes those POD5 files to generate TSV files that output f5c methylation calling. *Realfreq-program* calculates the modification frequencies. The absolute path to the custom pipeline should be given using the -s option (A sample f5c pipeline can be found in the scripts/pipeline-f5c.sh).

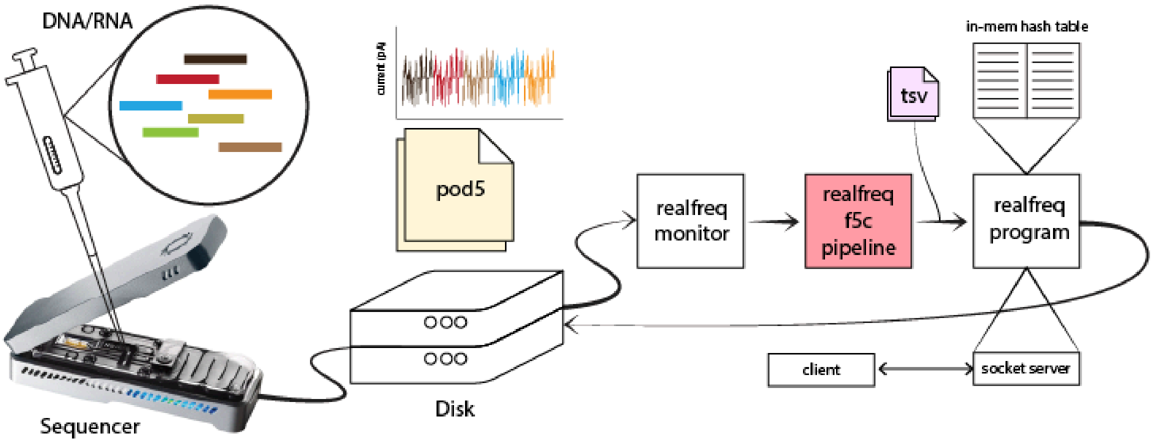


```
export DORADO_BIN=/ont-dorado-server/bin
export REF=/ref/hg38noAlt.fa
export REFIDX=/ref/hg38noAlt.idx
export DORADO_MODEL="dna_r10.4.1_e8.2_400bps_5khz_modbases_5hmc_5mc_cg_hac.cfg"
./scripts/realfreq.sh -m /data/exp_id -s scripts/pipeline-f5c.sh --tsv
```

#### Live modification calling

When ONT MinKNOW live modification calling is enabled ( by turning "Modified bases" and "Alignment" ON), as shown in the figure below, *realfreq-monitor* monitors modBAM files generated by ONT MinKNOW. The *realfreq-modbam-pipeline* (light yellow block in the figure) generated the index files that are then used by *realfreq-program* to calculate the modification frequencies. The file format (.bam) to monitor should be specified using the -a option. The absolute path to the custom pipeline should be given using the -s option (scripts/pipeline-modbam.sh).

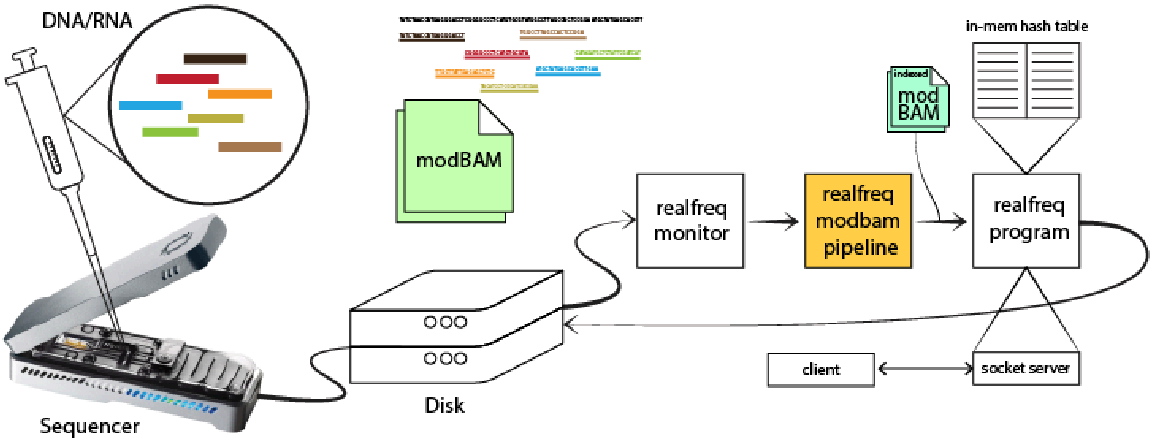


```
export REF=/ref/hg38noAlt.fa
./scripts/realfreq.sh -m /data/exp_id -s scripts/pipeline-modbam.sh -a "bam"
```

#### Command and options

##### Command

```
reafreq.sh -m [directory] [options ...]
```

##### Environment variables

Required for tools in pipeline scripts

- DORADO_BIN: Path to the directory containing Dorado binary
- REFIDX: Reference genome index for minimap2 alignment
  We need to provide a REFIDX (reference index) when using the default pipeline as minimap2 will not perform the repetitive computation of the index per each input file. The index can be generated from reference.fasta using Minimap2 using the command below
  ```
  minimap2 -ax map-ont reference.fasta -d reference.idx
  ```
- DORADO_MODEL: Basecalling + base modification model

Required for realfreq-script and realfreq-program

- REF: Reference genome
- REALFREQ_AUTO: If set to 1, realfreq.sh terminates automatically at the end of MinKNOW sequencing run.
- REALFREQ_THREADS: Number of threads used for modification calling (default 1)

##### Available options

```
OPTIONS
    -h, --help         Print help message
    -i, --info         Print script information
    -m [directory]     The sequencing experiment directory to be monitored
    -o [output]        Output file for modification frequency [default: freq.tsv]
    -r                 Resumes a previous live conversion
    -c [port]          Server port for realfreq
    -t [time]          Timeout in seconds [default: 21600]
```

```
    -p [processes]    Maximum number of parallel conversion processes [default: 1]
    -a [extension]    Watch for files with extension [default: pod5]
    -b                Output bedmethyl format
    -w INT            Write output (tsv/bedmethyl) every INT seconds (-1: only at
the end, 0: per batch) [default: 0]
```

### Supplementary Note 2 - Usage of the server-client configuration

*Realfreq-server* provides an interface to access the real-time modification frequency information using TCP socket connections. To launch this server,

*realfreq.sh* pipeline and realfreq don't start it unless specified by setting -c <port> or --server <port> flag. The server supports several query commands.

If the server starts up successfully, the following INFO log with the port number should be printed onto the terminal.

```
[main::INFO] Starting server on port 5001

[start_realfreq::INFO] reading file path from stdin
[start_server::INFO] server listening on port 5001.
```

##### Example querying commands demonstrated using netcat

```
nc localhost 8080 <<< help
nc localhost 8080 <<< get_contig:chr1
nc localhost 8080 <<< get_range:1:100
nc localhost 8080 <<< get_contig_range:chr22:18850302:49514860
nc localhost 8080 <<< get_contig_range_mod:chr22:18850302:49514860:m
```

##### Available query commands

```
help
    show this help message
get_contig:<contig>
    query by contig
get_range:<start_pos>:<end_pos>
    query data between start and end positions (both inclusive)
get_contig_range:<contig>:<start_pos>:<end_pos>
    query by contig and between start and end positions
get_contig_range_mod:<contig>:<start_pos>:<end_pos>:<mod_code>
    query by contig and between start and end positions and by mod code
```

### Supplementary Note 3 - Usage of realfreq-program as a standalone tool

Realfreq can be used standalone as a tool to compute live modification frequency. Realfreq reads the input modBAM file path from standard input. Therefore, the following two methods can integrate realfreq into an existing pipeline.

```
# using pipe
echo /path/to/reads.bam | ./realfreq -r ref.fa -o freq.tsv

# input a list of bam files
./realfreq -r ref.fa -o freq.tsv < bams_list.txt
```

Further, realfreq supports f5c/nanopolish modification calling TSV as input to compute modification frequencies.

Usage:

```
realfreq [options..] ref.fa
```

Options:

```
   -b                          output in bedMethyl format [not set]
   -c STR                      modification codes (ex. m , h or mh) [m]
   -m FLOAT                    min modification threshold(s). Comma separated values for
each modification code given in -c [0.8]
   -t INT                      number of processing threads [8]
   -K INT                      batch size (max number of reads loaded at once) [512]
   -B FLOAT[K/M/G]             max number of bytes loaded at once [20.0M]
   -h                          help
   -p INT                      print progress every INT seconds (0: per batch) [0]
   -o FILE                     output file [stdout]
   -d FILE                     dump file [realfreq.dump]
   -l FILE                     progress log file [realfreq.log]
   -r                          resume from dump file [no]
   -s PORT                     start server on PORT [-1]
   -v INT                      verbosity level [4]
   -V                          print version
   -w INT                      write output every INT seconds if new modifications found
(-1: only at the end, 0: per input) [0]
```

### Supplementary Note 4 - The commands used for evaluation

##### MinKNOW output batch size configuration

Recent MinKNOW versions output POD5 files at a fixed time interval (e.g., every 1 hour). We changed the MinKNOW configuration to write pod5 files with 4000 reads per file, how it used to be in older MinKNOW versions. This was done by editing the */opt/ont/minknow/conf/package/shared/default_writer.toml* file as below (see text in bold).

```
[writer_configuration.read_pod5] # pod5
disable_writing_passed_reads = false
disable_writing_failed_reads = false
disable_writing_force_skipped_reads = false

file_pattern =
"pod5{basecall_status}/{alias}/{flow_cell_id}{basecall_status}_{alias_}{short_protocol_run_id}_{short_run_id}_{batch_number}.pod5"

batch_duration = "0s"
batch_count = 4000
#no_output_based_batching = {} # empty dict converts to protobuf.Empty and signals the option is enabled

enable = []
```

##### Evaluating the default realfreq-pipeline

How *realfreq.sh* was executed to perform MinION evaluation for the default *realfreq-pipeline* on the laptop:

```
export REALFREQ_AUTO=1
export DORADO_BIN=/ont-dorado-server/bin
export REF=/ref/hg38noAlt.fa
export REFIDX=/ref/hg38noAlt.idx
export DORADO_MODEL="dna_r10.4.1_e8.2_400bps_5khz_modbases_5hmc_5mc_cg_hac.cfg"
./scripts/realfreq.sh -m /data/exp_id
```

The commands in the default pipeline (simplified) had the following parameters and arguments:

```
blue-crab p2s -p1 batch.pod5 -o batch.blow5
buttery-eel -g $DORADO_BIN --port 5000 --use_tcp --device cuda:all --call_mods --config dna_r10.4.1_e8.2_400bps_5khz_modbases_5hmc_5mc_cg_hac.cfg -i batch.blow5 -o unaligned.sam
samtools fastq -@8 -TMM,ML unaligned.sam > batch.fastq
minimap2 -t 8 -ax map-ont --sam-hit-only -Y -y --secondary=no /ref/hg38noAlt.idx batch.fastq > unsorted.sam
samtools sort -@ 8 -o sorted.bam unsorted.sam
samtools index -@ 8 sorted.bam
```

Tool versions:

- Blue-crab: 0.1.0
- Buttery-eel: 0.5.0
- Dorado basecaller: 7.2.13
- Samtools: 1.18
- Minimap2: 2.26
- MinKNOW: 24.06.14

The same above commands were used for the PromethION 2 solo evaluation on the desktop using MinKNOW: 23.11.4.

**Evaluating the f5c pipeline**

How realfreq-script was executed during the MinION evaluation for the f5c pipeline on the laptop:

```
export REALFREQ_AUTO=1
export DORADO_BIN=/ont-dorado-server/bin
export REFIDX=/ref/hg38noAlt.idx
export DORADO_MODEL="dna_r10.4.1_e8.2_400bps_5khz_modbases_5hmc_5mc_cg_hac.cfg"
./scripts/realfreq.sh -m /data/exp_id -s /realfreq/scripts/pipeline-f5c.sh
```

The commands in the f5c pipeline (simplified) had the following parameters and arguments:

```
blue-crab p2s -p1 batch.pod5 -o batch.blow5
buttery-eel -g $DORADO_BIN --port 5000 --use_tcp --device cuda:all  --config
dna_r10.4.1_e8.2_400bps_5khz_hac.cfg -i batch.blow5 -o batch.fastq
minimap2 -t 8 -ax map-ont --sam-hit-only --secondary=no /ref/hg38noAlt.idx batch.fastq >
unsorted.sam
samtools sort -@ 8 -o sorted.bam unsorted.sam
samtools index -@ 8 sorted.bam
f5c index --slow5 batch.blow5 batch.fastq
f5c call-methylation --slow5 batch.blow5 -b batch.bam -g /ref/hg38noAlt.fa -r batch.fastq >
batch.tsv
```

Tool versions:

- Blue-crab: 0.1.0
- Buttery-eel: 0.5.0
- Dorado basecaller: 7.2.13
- Samtools: 1.18
- Minimap2: 2.26
- f5c: 1.3
- MinKNOW: 24.06.14

**Evaluating the RNA pipeline**

How realfreq-script was executed during the MinION evaluation for the RNA pipeline on the PC:

```
export REALFREQ_AUTO=1
export DORADO_BIN=/ont-dorado-server/bin
export REF=/genome/gencode.v40.transcripts.fa
export REFIDX=/genome/gencode.v40.transcripts.idx
export DORADO_MODEL="rna_rp4_130bps_modbases_m6a_drach_sup.cfg"

realfreq_script=/realfreq/scripts/realfreq.sh
pipeline_script=/realfreq/scripts/pipeline-rna.sh

$realfreq_script -m /data/exp_id -s $pipeline_script
```

As realfreq-program is executed inside realfreq_prog() in realfreq-script, realfreq-script was modified as below to provide a modification code argument -c for the realfreq-program so that it computes m6A modification frequencies.
scripts/realfreq.sh

```
realfreq_proc(){
    ..............
```

```
    ...............
    ${REALFREQ} -c a -t $REALFREQ_THREADS -d $DUMP_FILE -o $OUTPUT_FILE $REF -l
$TMP_FILE_PATH $server_port_flag $bedmethyl_output_flag $resume_flag 2> $REALFREQ_PROG_LOG
}
```
